# Recurrent Duplication of the REST–NOA1 Locus Reveals a Hotspot of Regulatory Evolution Across Even-Toed Ungulates

**DOI:** 10.64898/2026.06.22.733815

**Authors:** Rick E. Masonbrink, Sharu Paul Sharma, Viswanathan Satheesh, Aleksandra Badaczewska-Dawid, Sivanandan Chudalayandi, David Alt, Paola Boggiatto, Ellie Putz, Andrew J. Severin, Steven C. Olsen

## Abstract

Gene duplication is a major source of evolutionary novelty, yet recurrent duplication of the same genomic locus across independent lineages remains understudied. Using a chromosome-scale assembly and de novo annotation of the North American bison (*Bison bison*), we investigated the mechanisms underlying immune-system diversification in ungulates. Comparative genomics of bison, cattle, elk, and 13 mammalian species identified a large segmental duplication encompassing the regulatory genes REST and NOA1. Although similar duplications were present in 14/16 species examined, phylogenetic analyses revealed that duplicated genes cluster within species rather than across taxa, indicating repeated independent origins rather inheritance from a common ancestor. To characterize the evolutionary consequences of this duplication, we integrated transcriptomics from six tissues with weighted gene co-expression and comparative protein clustering. We identified extensive structural divergence among duplicated copies of REST and NOA1, including domain losses, truncations, and expression partitioning across tissues, consistent with functional divergence following duplication. Comparative analyses revealed substantial remodeling of immune-related genes in bison, including BOLA class I molecules, T-cell receptor genes, leukocyte immunoglobulin-like receptors, and regulators of immune signaling. These modifications included domain rearrangements, novel sequence insertions, and lineage-specific duplications associated with immune-related expression networks. Together, these findings identify the REST-NOA1 locus as a recurrent hotspot of segmental duplication in ungulates and demonstrate how repeated duplication, structural divergence, and regulatory partitioning contribute to genome evolution. These results suggest that immune system diversification in bison and related species has been shaped not only by sequence evolution but also by recurrent remodeling of regulatory pathways and antigen-recognition pathways.

## Introduction

Gene duplication is a major source of evolutionary innovation, generating new genetic material that is suitable for modification through subfunctionalization, neofunctionalization, or changes in gene dosage(Ohno 1970b; Birchler and Yang 2022; Kuzmin et al. 2022). While many duplications arise once and are inherited through descent, some genomic regions appear predisposed to repeated duplication across independent evolutionary lineages. Such recurrent events provide opportunities to identify genomic features that promote structural variation and to understand how duplication contributes to the diversification of regulatory and adaptive pathways. Despite the importance of gene duplication in genome evolution, examples of repeated duplication of the same genomic region arising independently across large mammalian clades remains poorly characterized (Sémon and Wolfe 2007; Maeso et al. 2012).

Gene duplication and divergence are fundamental mechanisms driving adaptive evolution in mammals, particularly in immune system evolution (Flajnik and Kasahara 2010; Kasahara 2010). In bison, emerging research suggests that modifications in key immune components, including histocompatibility molecules and T cell receptors (TCR), may contribute to immune response diversity (Mikko et al. 1997; Radwan et al. 2007). Strong selection has been found among cattle and river buffalo major histocompatibility complexes (MHC), indicating a diversification of antigen presentation and adaptive immunity (Ren et al. 2021). TCR diversity is generated by variable-diversity-joining (V(D)J) recombination, a process responsible for generating highly variable antigen recognition receptors, and variation in the TCRB locus has been linked to differential immune responses in mammals (Russell et al. 2022). Although bison experienced a severe population bottleneck that reduced overall genetic diversity, previous studies have identified substantial variation within immune-related loci, suggesting that adaptive immune genes may follow evolutionary trajectories distinct from genome-wide patterns (Mikko et al. 1997; Radwan et al. 2007). Comparative genomic analyses are thus invaluable or revealing how adaptation and divergence have shaped immune responses, as even closely related species like cattle and goat exhibit massive differences in TCR gene structure and usage (Gillespie et al. 2021). Understanding these genetic modifications in bison can provide insight into broader patterns of immune system evolution among even-toed ungulates, revealing how gene duplication and divergence have shaped immune defense strategies over evolutionary timescales (Rodriguez et al. 2022).

Regulatory genes are particularly attractive targets for duplication because changes in dosage or expression can have cascading effects on downstream gene networks. Recent comparative genomic studies have revealed that some genomic regions repeatedly undergo structural rearrangement in multiple lineages, suggesting the existence of genomic hotspots for duplication and diversification (Maeso et al. 2012; Sharma and Peterson 2023). Such regions are of particular interest as they provide insight into the mechanisms generating genome novelty and evolutionary processes that maintain duplicated genes. Determining whether similar duplications observed across species arise from a shared ancestral event or through independent convergence remains a central challenge in understanding genome evolution (A. von der Dunk and Snel 2020).

Even-toed ungulates provide an informative system for investigating the evolutionary consequences of gene duplication because they occupy diverse ecological niches and experience a wide range of pathogen pressures. Comparative genomic studies have revealed extensive diversification of immune-related loci among ruminants, including histocompatibility complexes, immunoglobulin genes, and T cell receptor loci (Gillespie et al. 2021; Masonbrink et al. 2021; Oppenheimer et al. 2021b; Ren et al. 2021). Despite this growing understanding of sequence-level divergence, the contribution of recurrent structural variation and segmental duplication to immune system diversification across ungulates remains largely unexplored.

Bison represent a particularly informative system for examining these processes. In addition to experiencing severe demographic bottlenecks and some limited introgression from domestic cattle, bison occupy a unique position among ruminants as an ecologically important wildlife species which also happens to be a reservoir for several livestock pathogens (Halbert and Derr 2007; Hedrick 2009; Stroupe et al. 2022). Previous studies have identified divergence in immune-related loci among bison, cattle, and elk, suggesting that lineage-specific modifications of immune pathways may contribute to differences in host responses (Mikko et al. 1997; Radwan et al. 2007; Ren et al. 2021). These characteristics make bison a useful model for investigating how structural genome evolution contributes to immune system diversification.

Here, we identify a recurrent segmental duplication encompassing the regulatory genes REST and NOA1 across multiple ungulate lineages. Using a chromosome-scale assembly and de novo annotation of the North American bison together with comparative genomics, transcriptomics, and phylogenetic analyses, we demonstrate the duplicated copies have diverged in both structure and expression and are associated with extensive remodeling of immune-related genes. These findings reveal a recurrent source of regulatory innovation in ungulate genomes and provide insight into the mechanisms by which duplication and divergence contribute to immune system evolution.

## Methods

### Genome sequencing and chromosomal assembly

High-molecular weight DNA was isolated from blood collected from two adult male Yellowstone bison. Long-read sequencing was performed on an Oxford Nanopore GridIion X5, generating 95.4Gb of sequence data from 8.75 million reads. Chromatin capture libraries were generated using the Phase Genomics Proximo Hi-C 3.0 kit and sequenced on an Illumina NovaSeq 6000, yielding 207.4 million paired end reads (Lieberman-Aiden et al. 2009).

Nanopore reads were assembled with FLYE and scaffolded using the Phase Genomics Proximo Hi-C pipeline followed by manual curation in Juicebox (Bickhart et al. 2017) (Supplementary Figure 1). The final assembly consisted of 31 chromosome-scale pseudomolecules and 7 additional scaffolds. Assembly quality was assessed using BUSCO, yielding completeness scores of 98.1% for eukaryote_odb10, 89.0% for mammalia_odb10, and 85.8% for cetartiodactyla_odb10 (Supplementary Figure 2). Additional assembly, polishing, and quality control procedures are described in the supplementary methods.

**Supplemental Figure 1.**
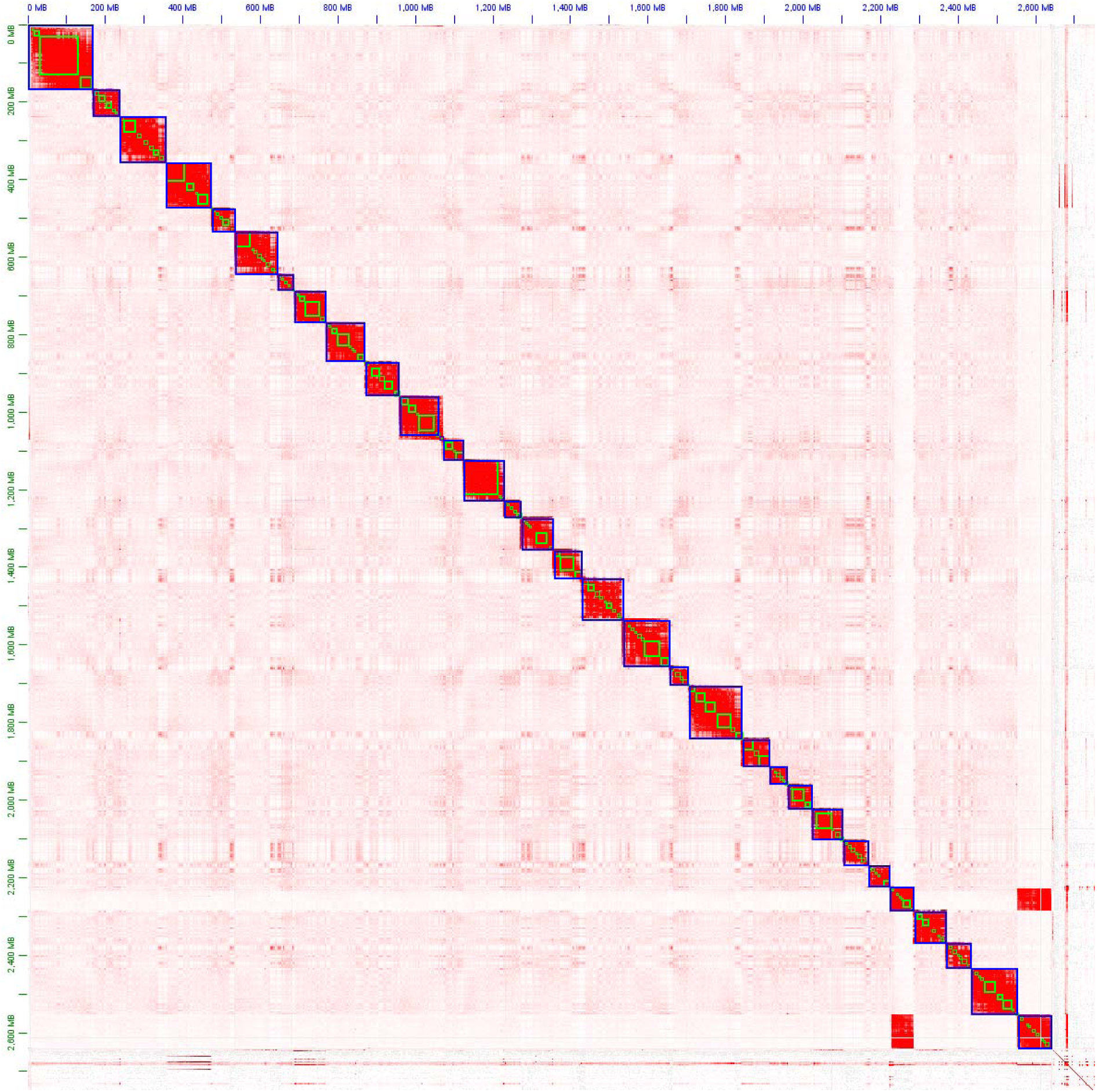
Hi-C reads aligned to the *B. bison* genome representing the final 31 pseudomolecule scaffolds visualized in JuiceBox. Cross homology of the last scaffold represents homology between the X and Y chromosomes.

**Supplemental Figure 2.**
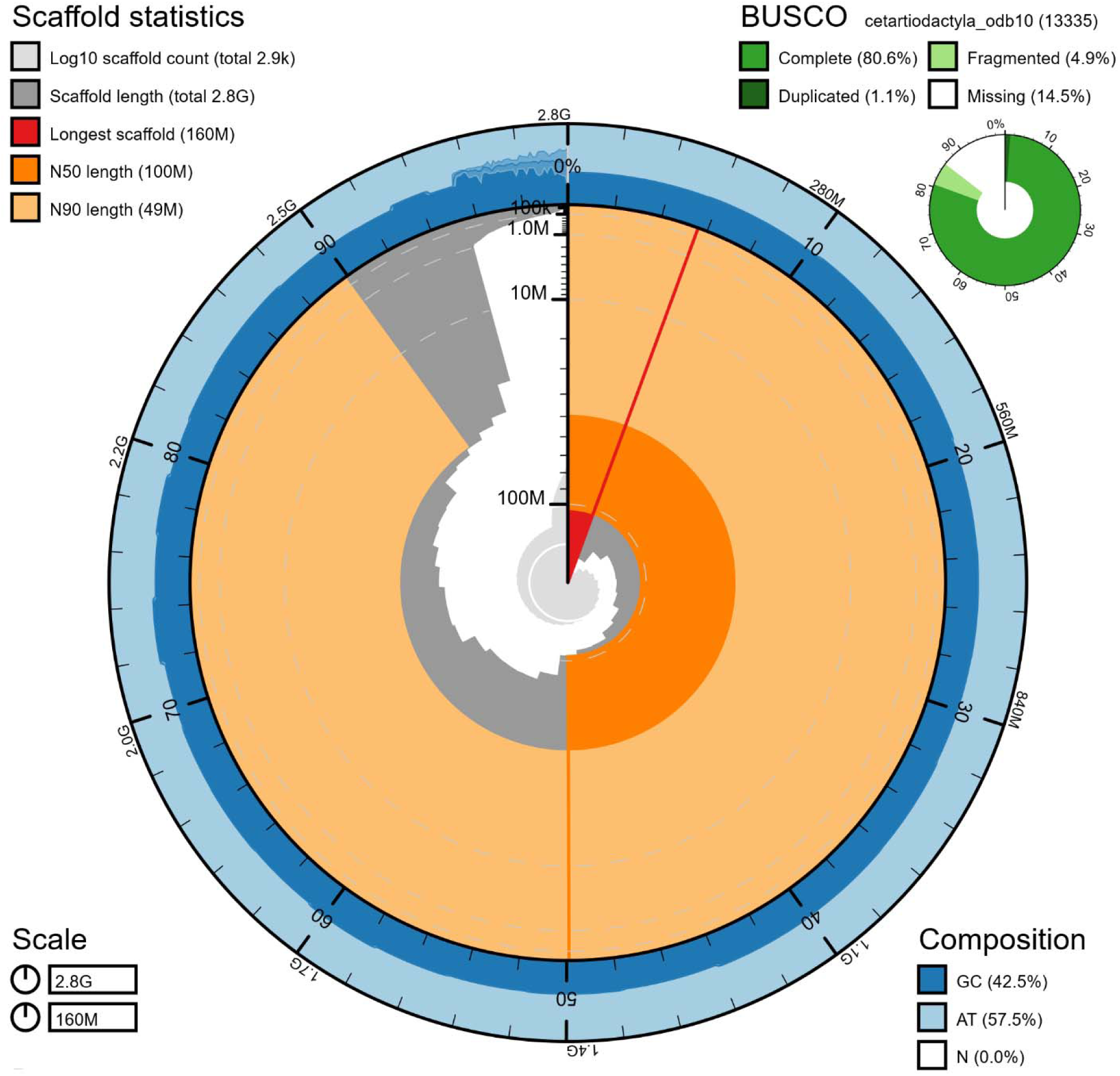
Snail plot represents the genomic and BUSCO statistics of this bison assembly.

### Gene prediction and functional annotation

Genome annotation was performed using a consensus transcript approach implemented in Mikado (Mapleson et al. 2018; Venturini et al. 2018). Repetitive elements were identified and soft-masked prior to annotation (Smit et al. 2014), and transcript evidence was generated from multiple RNA-seq assembly methods (Haas et al. 2013; Pertea et al. 2016; Song et al. 2016; Liu and Dickerson 2017; Hoff et al. 2018; Mao et al. 2020) together with protein homology from bison, cattle and elk (Gremme 2014). Functional annotation was assigned using DIAMOND searches against NCBI NR and UniProt databases (Apweiler et al. 2004; Suzek et al. 2007; Buchfink et al. 2015; Consortium 2017; Consortium 2019). Software versions and additional implementation details are provided in Supplementary Table 1 and the Supplementary Methods.

### Comparative genomic analyses

The predicted proteomes of bison, elk, and cattle were clustered with the current assembly’s predicted proteins using Orthofinder (Emms and Kelly 2015). Orthologous annotations were obtained using Diamond (Buchfink et al. 2015) to BLAST (Madden 2013) each predicted protein to Uniprot (Apweiler et al. 2004; Suzek et al. 2007; Consortium 2017; Consortium 2019).

We assessed synteny with ntSynt (Coombe et al. 2025a) and ntSynt-viz (Coombe et al. 2025b) amongst bison, elk, and cattle genome alignments with minimap2 (Li 2016). Genomes used for gene duplication analysis are listed in Supplementary Table S2.

### REST-NOA1 duplication analyses

First, we used Genomethreader (Gremme 2014) alignments of the four duplicated bison genes to identify the locus in the genome. We then extracted these regions and 500kb surrounding them for processing with BLAST (Madden 2013) self-alignments. Using BLAST dotplots, approximate locations of each duplications’ copy were identified. These duplications were visualized in JBROWSE2 (Diesh et al. 2023), along with the Genomethreader (Gremme 2014) alignments and the NCBI Refseq gene annotations (O’Leary et al. 2016) for each species when available. To identify repetitive elements, we used RepeatModeler and RepeatMasker on selected chromosomes from each species.

To account for variation in annotation software, we extracted 1Mb regions surrounding this duplication and annotated the genes in this region with Augustus using human training. The protein and CDS sequences were extracted from REST and NOA1 genes with gffread and aligned using Mafft with – maxiterate 1000 –localpair –op 5 and –ep 0.1 (Katoh and Standley 2013). Trimal was used to trim poorly aligned regions. Maximum likelihood trees were built from these alignments in RAxML using automatic model selection with Gamma-distributed rate heterogeneity and 100 bootstraps. Trees were displayed in iTOL and rooted to *T. kanchil* (Letunic and Bork 2024). A species tree was created with Orthofinder (Buchfink et al. 2015; Emms and Kelly 2015).

### Transcriptomic analyses

Gene expression was assessed using STAR (Dobin et al. 2013) alignments of 165Gb of RNAseq from 6 tissues: (adult female kidney, adult female liver, adult female lung, adult female skeletal muscle, adult female supra-mammary lymph node, and adult female spleen), (SRR1659047-SRR1659070). Read counts and differential expression were obtained using featureCounts from the Subread package and Deseq2 (Liao et al. 2013; Love et al. 2014).

For unsupervised WGCNA analysis (Langfelder and Horvath 2008), we retained bison genes based on their expression variability across samples. Specifically, genes with a variance (count data) higher than the 25th percentile of all gene variances were selected for further analysis. This resulted in 16,711 genes meeting the criteria. The count data from this set of genes was DESeq2-normalized (rlog) and were subsequently used for network evaluation. The WGCNA package in R provided the blockwiseModules function which was employed to form a signed network based on a Pearson correlation matrix. The entire gene set was processed as a single unit, and only positive correlations were considered in this signed network. The soft power threshold was set to 2, which is the minimum required to attain a scale-free topology with R2⍰=⍰0.8. Modules were identified using standard parameters with a mergeCutHeight of 0.15 and PAMstage activated. The first principal component of the expression data for a module determined its module eigengene expression pattern. The intramodular connectivity (kME) for each gene was established based on the Pearson correlation between the gene and the module eigengenes.

## Results

### Genome assembly and annotation

A chromosome scale assembly of the North American bison genome was generated using Nanopore and Hi-C sequencing data. The final assembly consisted of 31 pseudomolecules and 7 unscaffolded contigs, exhibiting high completeness, with BUSCO scores of 98.1% and 85.8% for the eukaryote_odb10 and cetartiodactyla_odb10 datasets (Supplementary Figure 1). Genome-wide alignments with previously published bison assemblies demonstrated extensive structural concordance, supporting assembly accuracy (Supplementary Figure 3) (Oppenheimer et al. 2021a; Stroupe et al. 2023). Comparisons with cattle and elk revealed broad conservation of chromosome structure together with lineage-specific rearrangements (Figure 1).

**Figure 1.**
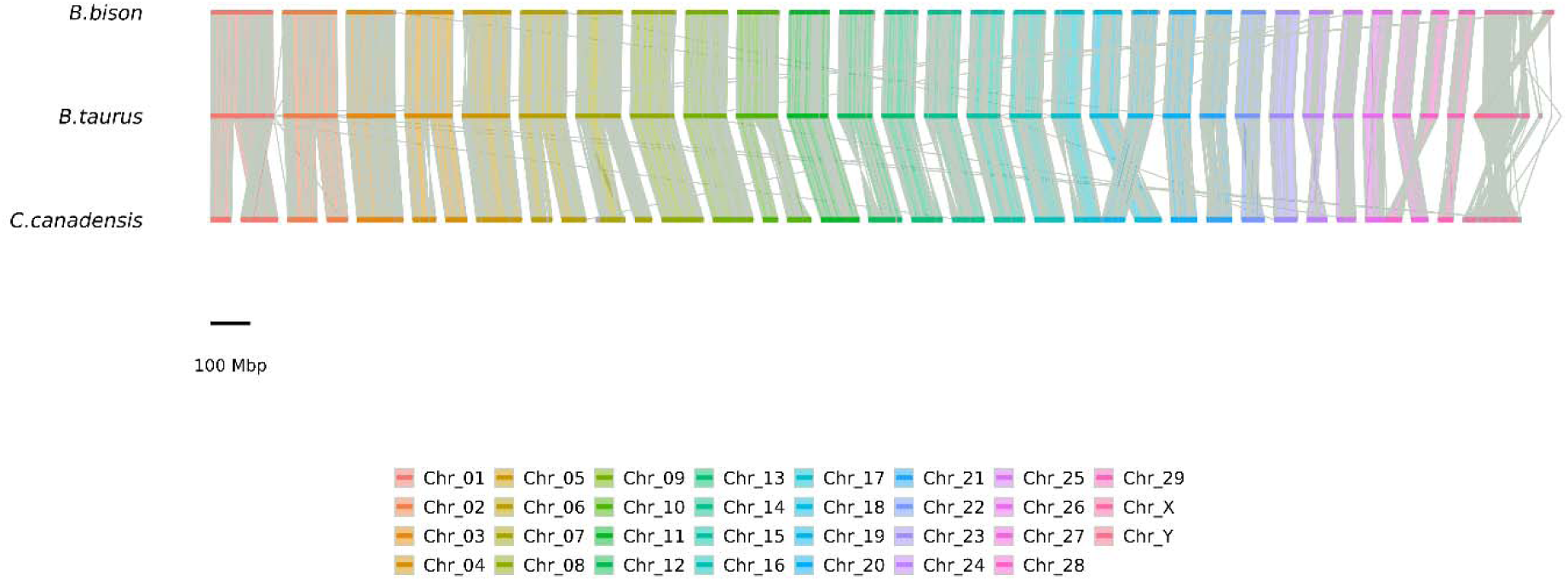
An ntSynt visualization of synteny between *B. bison*, *B. taurus*, and *C. canadensis*. Grey lines within chromosomes represent the borders between scaffolded contigs, while grey lines between chromosomes represent lower confidence unchained syntenic links. Synteny between *B. bison* and *B. taurus* was extensive and represented few rearrangements between chromosomes, excluding the end of chromosome 1 and the sex chromosomes. Synteny between B. taurus and C. canadensis was also extensive, with the same chromosome1 end translocations as found in B. bison, along with chromosome fissions/fusions and extensive rearrangements of sex chromosomes. Chromosome numbering corresponds to pseudomolecule labels in *B. bison*.

**Supplemental Figure 3.**
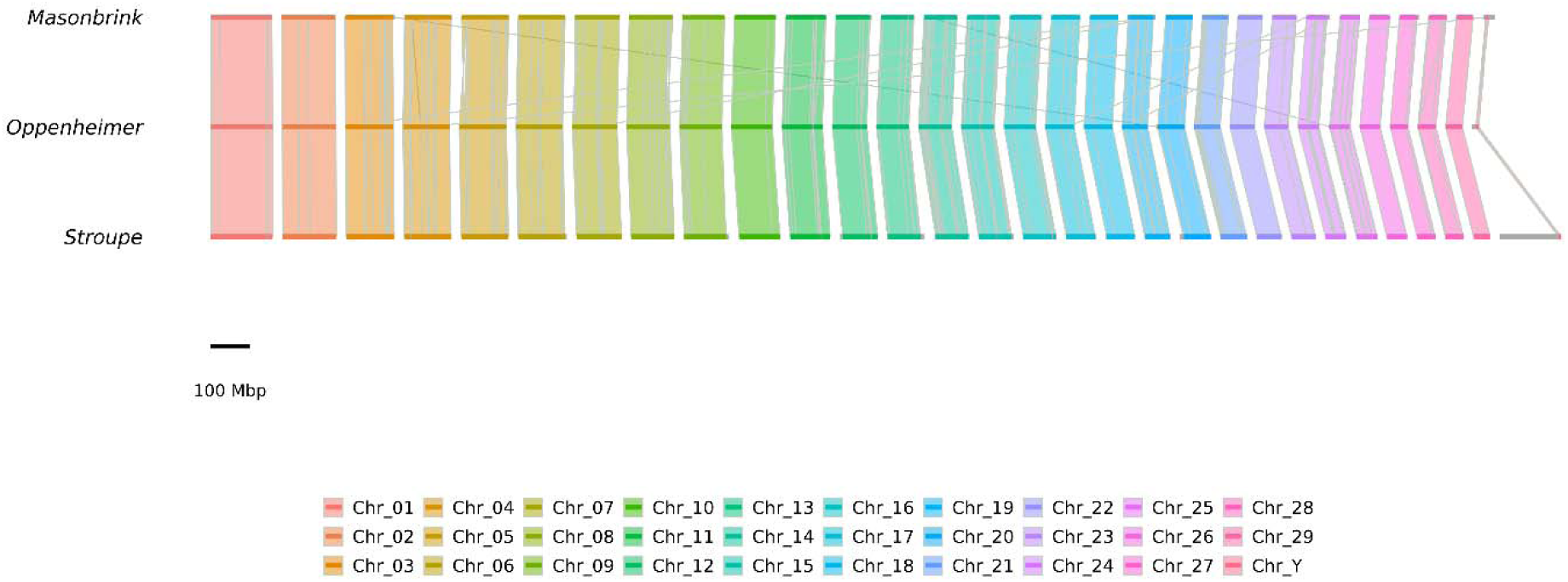
An ntSynt visualization of synteny between chromosomal genome assemblies of *B. bison*. Grey lines within chromosomes represent the borders between scaffolded contigs, while grey lines between chromosomes represent lower confidence unchained syntenic links. Chromosome numbering is consistent across assemblies, with some exception represented by homology between Y and X chromosomes.

Consensus-based gene prediction identified 22,282 transcripts representing 18,317 genes, including 3,402 genes with alternative splicing. Functional annotation was assigned to 98.3% of transcripts and all predicted genes, demonstrating extensive support from existing protein databases. Annotation completeness was high, with BUSCO scores of 98.8% and 93.8% for eukaryota_odb10 and certartiodactyla_odb10 datasets, respectively.

### Divergence of immune-related genes in bison

To identify candidate genes associated with immune-system diversification, we examined incongruencies within orthogroups to find lineage-specific structural modifications in bison relative to cattle and elk. Candidate genes were prioritized based on transcriptional support, immune-related functional annotation, and presence of substantial protein-level differences including domain gain or loss, internal duplication, truncation, or novel sequence insertions. This approach identified 12 immune-related loci exhibiting pronounced structural divergence (Table 1; Supplementary Table 3).

**Table 1.** Structurally divergence immune-related genes identified in bison relative to cattle and elk orthologs. Candidate genes were selected based on transcriptional support, immune-related functional annotation, and the presence of substantial protein differences, including domain gain or loss, internal duplication, truncation, tandem duplication, or novel sequence insertions. This table summarizes the major structural modifications identified in each gene and their associated functional categories.

| Gene ID | Annotation | Structural Modification | Functional Category |
| --- | --- | --- | --- |
| Bisbis_gene7931 | BOLA Class I antigen | Loss of N-terminal signal peptide | Antigen presentation |
| Bisbis_gene7967 | BOLA Class I antigen | Signal peptide replaced by disordered domain | Antigen presentation |
| Bisbis_gene7928 | BOLA Class I antigen | Novel 170-aa N-terminal extension | Antigen presentation |
| Bisbis_gene13119 | T-cell receptor beta chain | Domain loss, loss of transmembrane region in one isoform | T-cell recognition |
| Bisbis_gene18001 | T-cell receptor alpha chain | Additional 38-aa C-terminal extension | T-cell recognition |
| Bisbis_gene18263 | T-cell receptor alpha-like gene | 2.5-fold internal duplication and acquisition of additional IG domains | T-cell recognition |
| Bisbis_gene18274 | T-cell receptor alpha-like gene | Tandem duplication<br>resulting in expanded<br>IG domains | T-cell recognition |
| Bisbis_gene11525 | Rap1A | Loss of one PR00449<br>Ras-signature domain | Immune signaling |
| Bisbis_gene5707 | Leukocyte immunoglobulin-like<br>receptor | Novel 43-aa N-terminal<br>extension | Immune signaling |
| Bisbis_gene3114 | Lambda-1 light chain | Transcript<br>diversification and<br>novel immunoglobulin-like domains | Antibody diversity |
| Bisbis_gene13343 | hnRNP A2/B1 | 60-aa insertion and<br>RNA-binding domain<br>truncation | RNA processing |
| Bisbis_gene14091 | REST | Loss of DNA-binding<br>and REST-B domains | Transcriptional<br>regulation |

Comparative clustering of predicted proteins from bison, cattle, and elk identified 20,974 orthogroups, including 14,050 shared among all assemblies, indicating extensive conservation of protein-coding genes across these species. Despite this conservation, a mall subset of immune-related genes exhibited substantial structural divergence. These genes included members of the bovine leukocyte antigen (BOLA) family, T-cell receptor loci, immunoglobulin genes, and additional regulators of immune signaling and transcriptional regulation (Table 1).

The most pronounced differences were observed among BOLA class I genes, T-cell receptor loci, and the transcriptional regulator REST. Three BOLA class I genes exhibited modifications affecting N-terminal architecture, including loss of canonical signal peptides, replacement of signal peptides with disordered regions, and acquisition of a novel 170 amino-acid extension. Several T-cell receptor genes displayed domain loss, internal duplication, tandem duplication, and expansion of immunoglobulin-like domains relative to cattle and elk orthologs. Additional divergence was observed among genes involved in immune signaling, immunoglobulin structure, and RNA processing, including Rap1A, leukocyte immunoglobulin-like receptors, immunoglobulin lambda chains and hnRNP A2/B1 (Table 1; Supplementary Table 3).

Among the divergent genes identified, REST was unique in occurring within a large segmental duplication that also encompassed nitric oxide associated 1 (NOA1). Comparison across three independent bison assemblies confirmed the duplication and excluded assembly artifact as an explanation. Because structural variation affecting transcriptional regulators has the potential to alter downstream gene networks, this observation motivated a broader investigation of the locus across ungulates.

### Recurrent independent duplication of the REST-NOA1 locus across even-toed ungulates

Bisbis_gene14091 is located within a large segmental duplication that has created two additional genes: REST and NOA1. This duplication was confirmed by comparing three bison assemblies, supporting its biological existence. Comparative analysis of 16 mammalian genomes identified similar duplications in 14 species, suggesting that this genomic region represents a recurrent target of structural evolution in ungulates. Although the duplicated regions shared a conserved core gene set centered on REST and NOA1, duplication size, sequence identity, and affected genes varied substantially among lineages (Table 2; Supplementary Table 4).

**Table 2.**
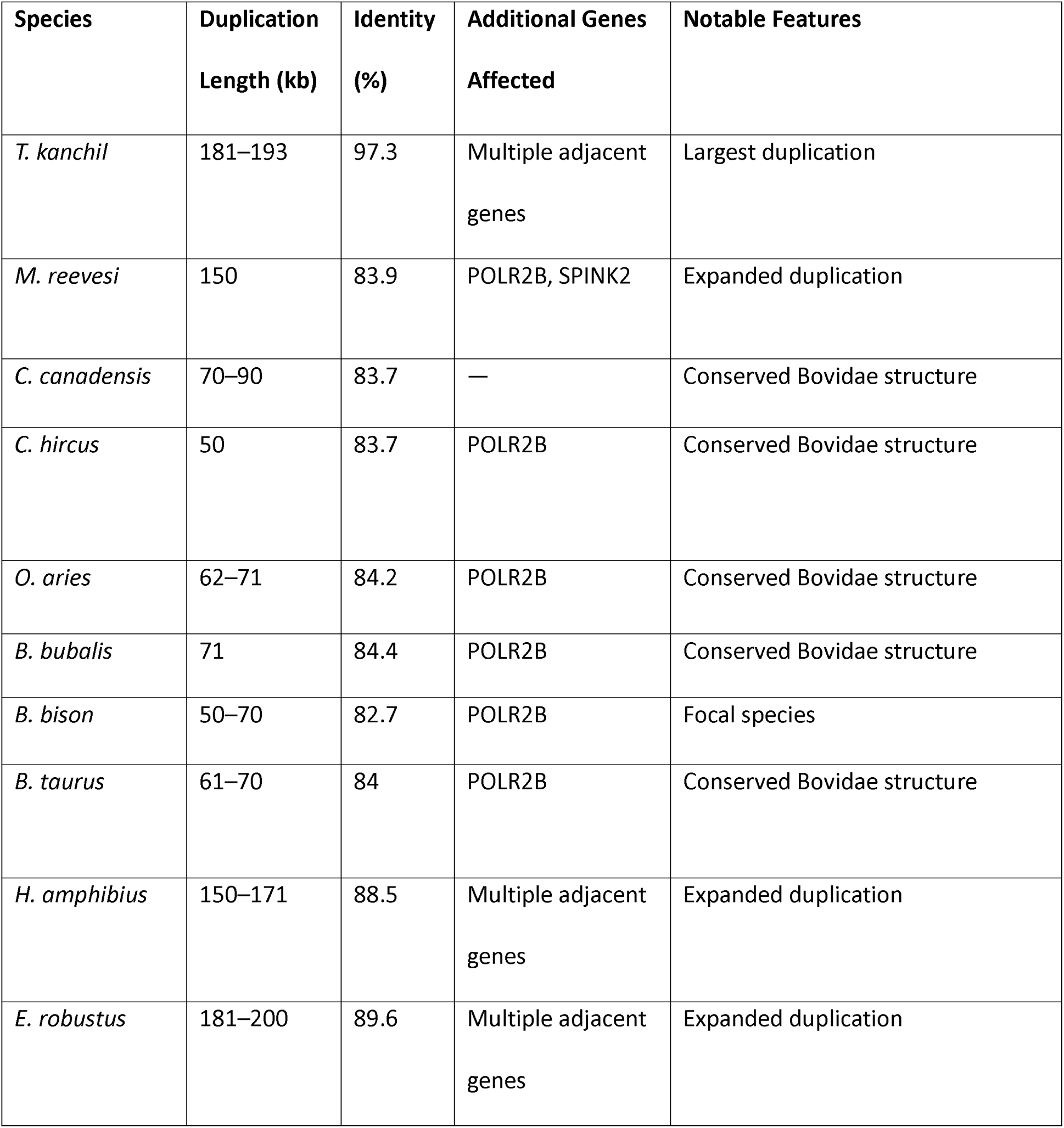
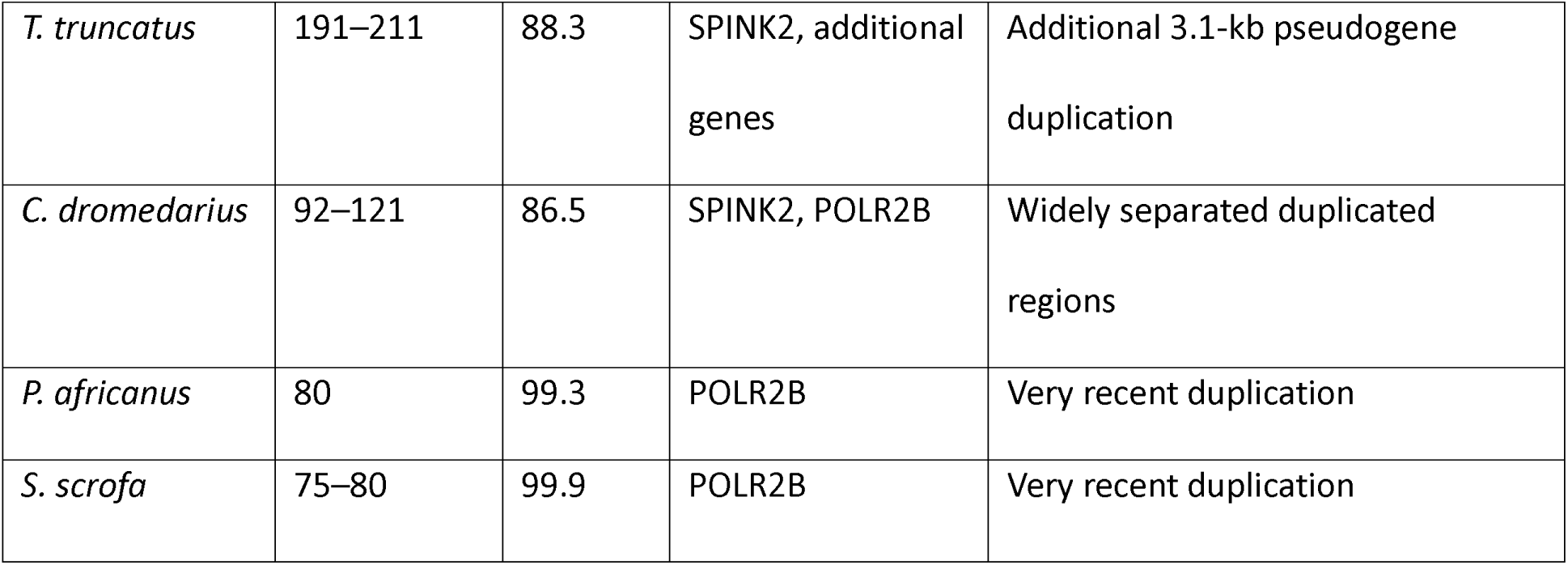
Recurrent duplication of the REST–NOA1 locus across ungulates. Duplication size, sequence identity, and affected genes are summarized for each species. REST and NOA1 were involved in all detected duplications except in *Equus asinus* and *Catagonus wagneri*, the only species in which no duplication was identified. Additional lineage-specific genes, expansions, pseudogenization events, and structural modifications illustrate substantial divergence following duplication.

Maximum likelihood phylogenies of REST CDS and proteins recovered duplicated copies clustering within species rather than by duplicate class, supporting multiple independent duplication events (Figure 2). NOA1B genes, often fragmented, did not show this pattern clearly, but variation in duplication size and sequence identity supports independent origins (Supplemental Figure 4). Duplication size and affected gene content varied by lineage (Supplementary Table 4), with some species having larger duplications affecting additional genes. Analysis of flanking regions showed substantial homology and high concentrations of repeats (e.g. RTE-BovB, SINE/tRNA), implicating their involvement in genomic rearrangement (Supplementary Table 5; Supplementary Table 6; Supplementary Table 7) (Sharma et al. 2021; Sharma and Peterson 2023). Several lineages also exhibited unique structural modifications of the locus. For example, *Camelus dromedarius* possessed an unusually large interval separating duplicated regions, whereas Tursiops truncatus retained an additional 3.1kb duplication on a separate chromosome containing RESTC and NOA1 pseudogene, further illustrating the dynamic evolutionary history of this genomic region.

**Figure 2.**
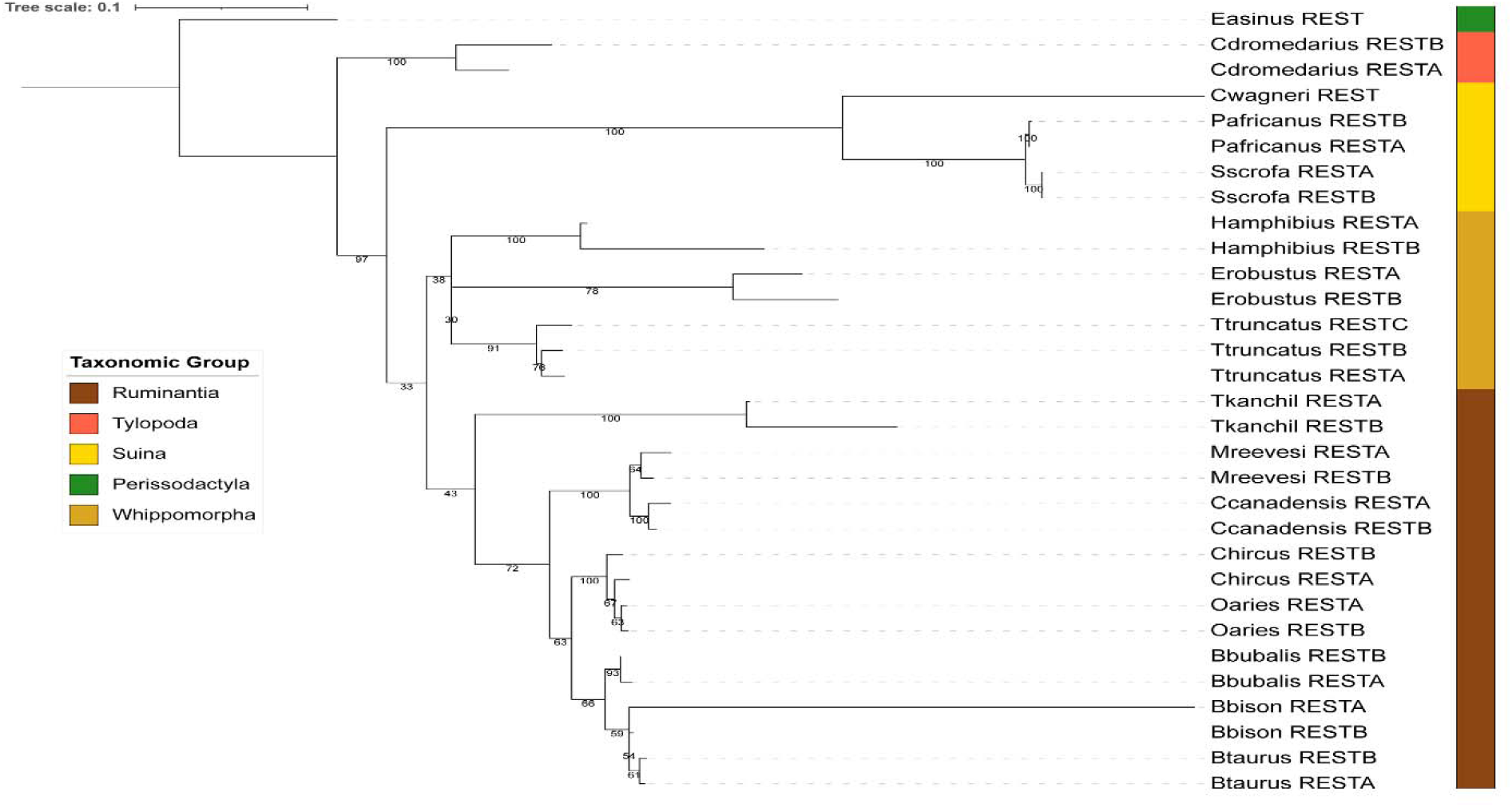
This is a maximum likelihood phylogenetic tree based on protein alignments of RESTA and RESTB. The REST genes found in the segmentally duplicated and were named according to their synteny between the species. Only one REST gene was found in *C. wagneri* and *E. asinus*, species that lacked this duplication. Numbers represent bootstrap support.

**Supplemental Figure 4.**
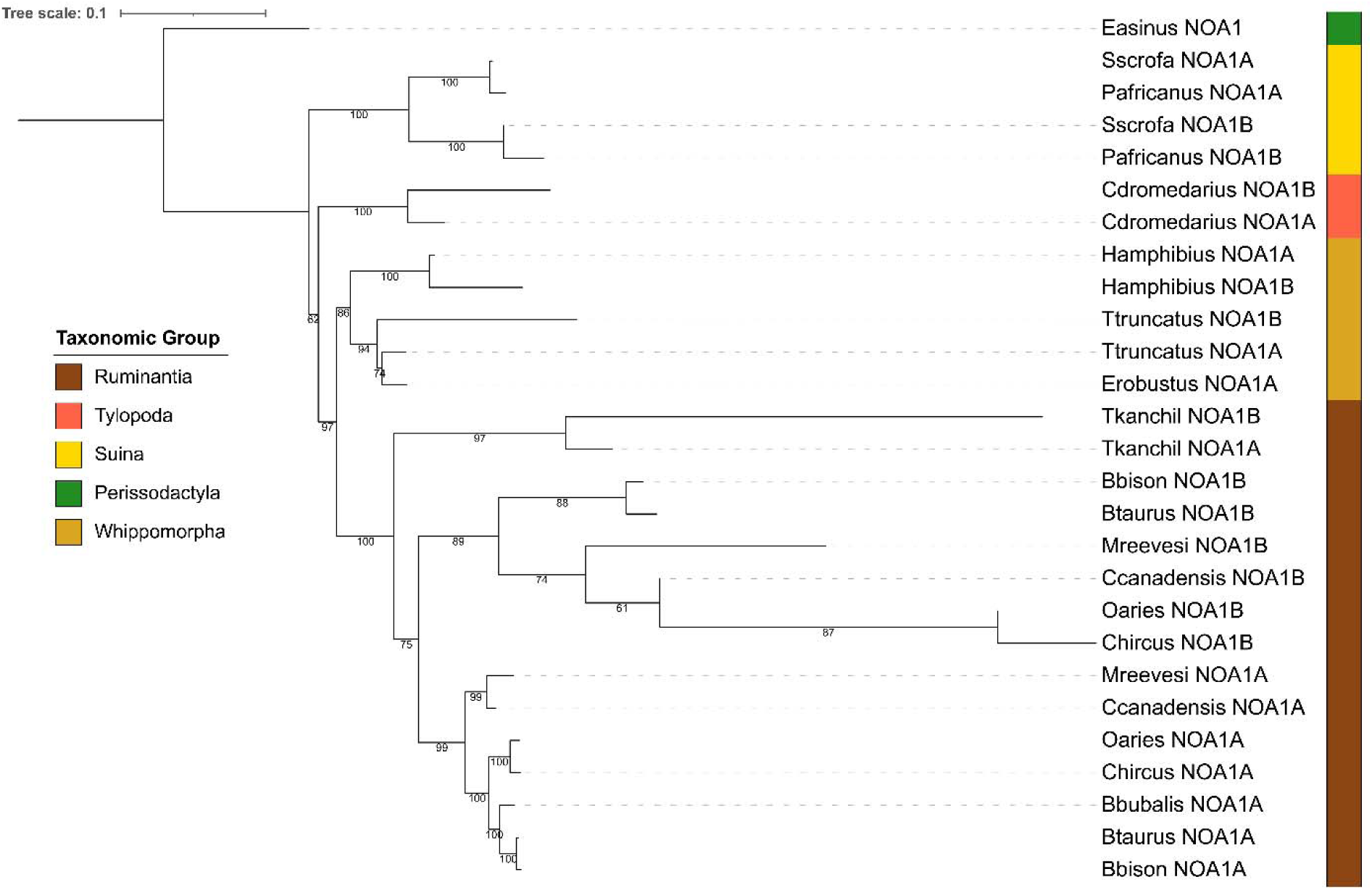
Maximum likelihood phylogenetic tree of CDS sequences illustrating the presence of duplicated NOA1 genes among 16 species. While many NOA1A and NOA1B genes cluster separately, this is due to the highly truncated nature of NOA1b genes, limiting their distinction in the tree. *E. asinus* and *C. wagneri* lacked the duplication and thus the duplicated genes. *B. bubalis* and *E. robustus* NOA1B genes became pseudogenized and are not represented in the tree. Numbers represent bootstrap support.

Duplicated copies of REST and NOA1 exhibited structural and expression divergence in bison. RESTA lacked a DNA binding domain present in RESTB, whereas RESTB lacked the RE-silencing transcription factor B domain retained in RESTA. Similarly, the YqeH GTPase domain of NOA1B was severely truncated relative to NOA1A. Expression profiles also differed substantially with RESTA showing elevated expression in immune tissues, RESTB exhibiting higher expression in liver and kidney, and NOA1A displaying broader expression than NOA1B. These duplicated copies differed substantially in both domain architecture and tissue-specific expression.

### Gene co-expression network analysis and immune system association

Weighted gene co-expression network analysis assigned 16,711 genes to 25 expression modules (Figure 3). Ontological enrichment identified three immune-associated modules enriched for inflammatory response, T cell mediated immunity, and antiviral defense pathways (Supplemental Figure 5). Several structurally divergent immune genes, including duplicated REST and NOA1 loci, were associated with these immune-related expression networks. RESTA was associated with immune-enriched modules whereas RESTB showed elevated expression in non-immune tissues.

**Figure 3.**
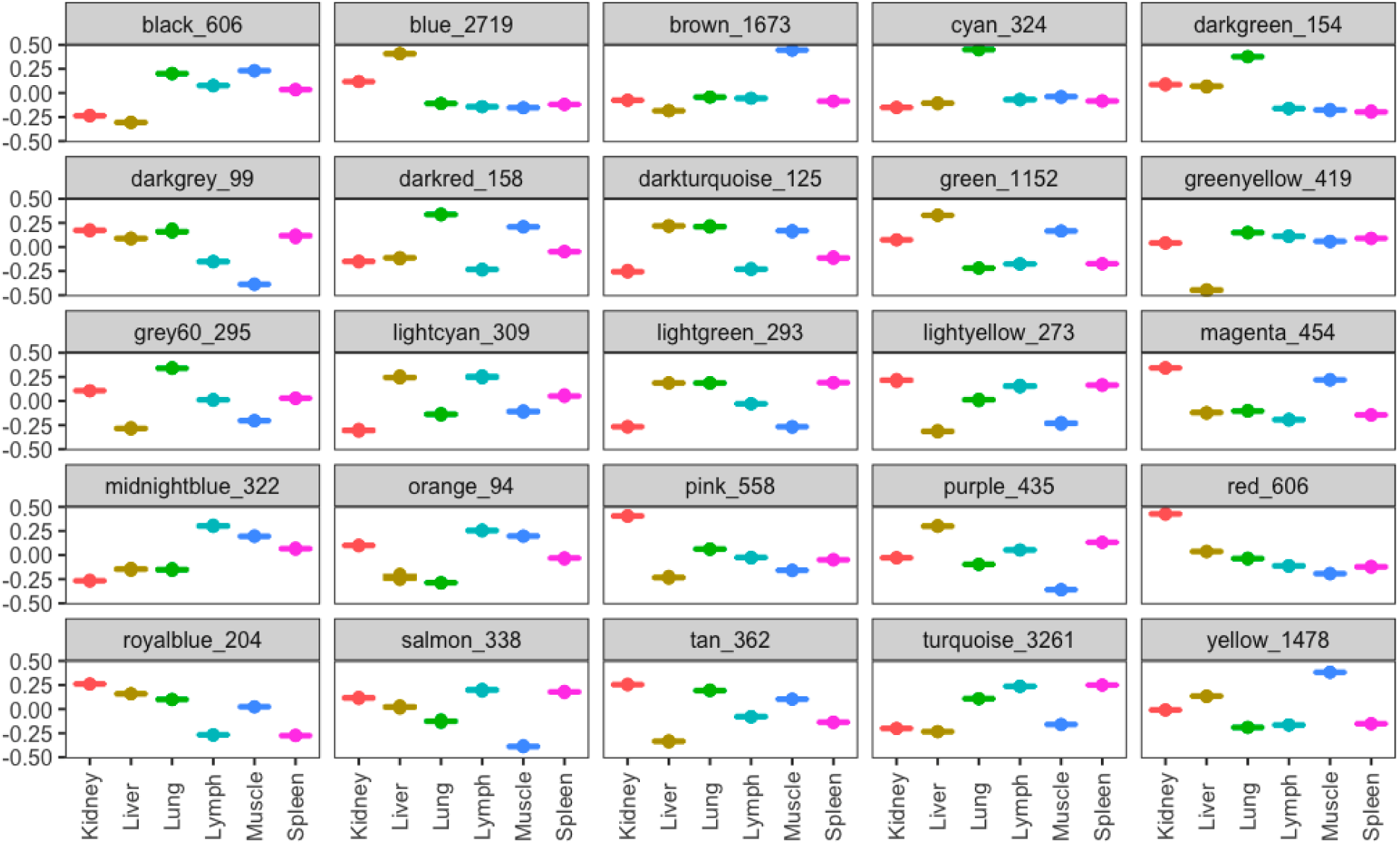
Distribution of Pearson correlation values across different tissues for various modules. Each subplot represents a module and contains box plots for the six tissues, allowing for comparison of distribution of Pearson correlations across multiple tissues. The numbers after each color represent the number of genes in each module.

**Supplemental Figure 5.**
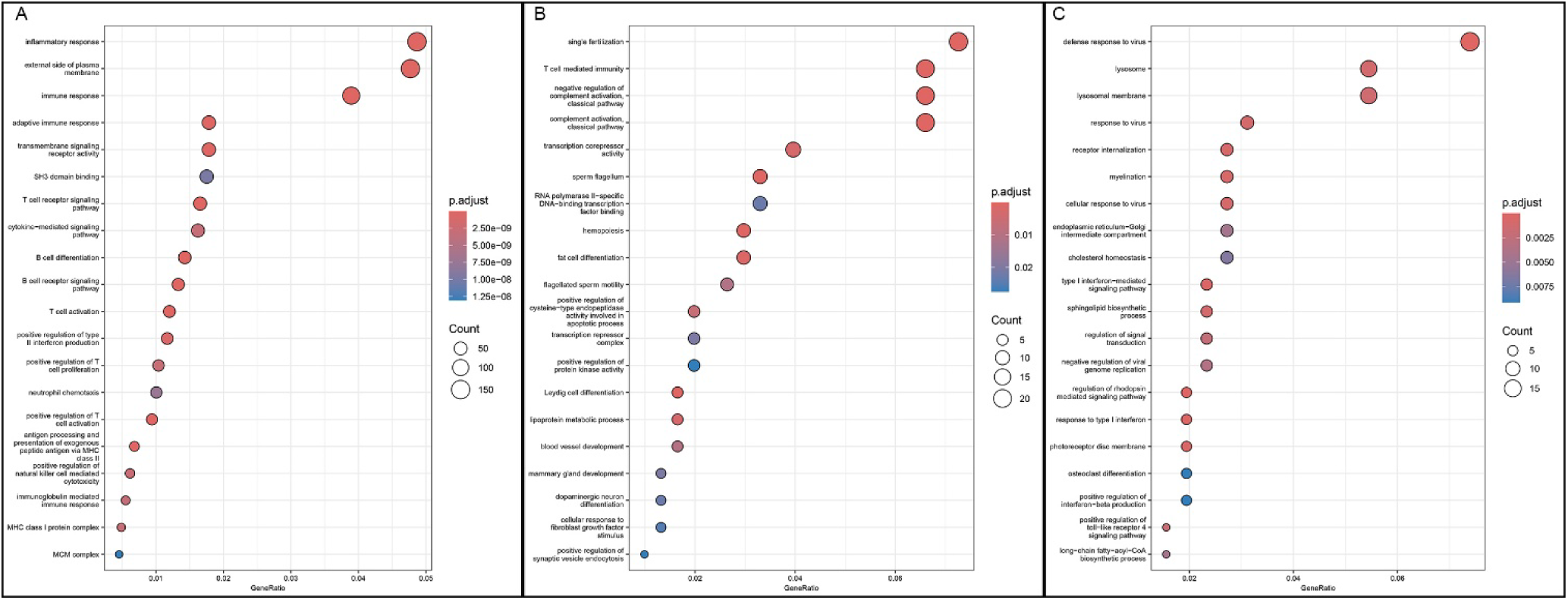
Clusterprofiler GO enrichments of twenty terms significantly enriched in a respective WGCNA expression module. The more significant a term, the further to the right its dot appears, while the size of the dot represents the number of genes with this GO term. A. represents the Turquoise module. B. represents the Midnight Blue module. C. represents the Light Yellow module.

## Discussion

Gene duplication is a major driver of genome evolution, yet examples of recurrent duplication affecting the same genomic region across independent mammalian lineages remain uncommon. Here, comparative analyses of bison and related ungulates identified repeated duplication of the REST-NOA1 locus together with extensive structural divergence among immune-related genes. These findings suggest that recurrent duplication, structural remodeling, and regulator divergence have contributed to immune-system evolution across even-toed ungulates and reveal the REST-NOA1 region as a previously unrecognized hotspot of genome evolution.

A central finding of this study is that similar REST-NOA1 duplications are present across multiple ungulate lineages but appear to have arisen independently rather than through inheritance from a common ancestral duplication. If these duplications originated from a single ancestral event, duplicated copies would be expected to cluster by duplicate class across species. Instead, duplicated REST genes consistently clustered within species, supporting repeated independent origins of the duplication. The repeated involvement of similar genes, together with extensive sequence homology and repeat-rich flanking regions, suggests that this genomic region is particularly susceptible to duplication through mechanisms such as non-allelic homologous recombination. These observations indicate that recurrent structural variation may represent an important and previously underappreciated source of genome innovation in even-toed ungulates. The repeat-rich sequence surrounding the locus likely facilitates recurrent duplication through mechanisms such as non-allelic homologous recombination mediated by transposable elements, specifically RTE-BovB and L1/SINE retroelements (Robberecht et al. 2013; Cerbin and Jiang 2018), whose distribution aligns with the two phylogenetic groupings observed.

Following duplication, REST and NOA1 copies exhibited substantial divergence in both domain architecture and tissue-specific expression. RESTA and RESTB retained different subsets of functional domains and displayed distinct expression profiles, while NOA1B exhibited substantial truncation relative to NOA1A. Such divergence is consistent with functional differentiation following duplication, a process that can partition ancestral functions or facilitate the evolution of novel regulatory roles. Because REST functions as a chromatin-associated transcriptional regulator with broad effects on downstream gene networks, structural modification of duplicated copies may have consequences extending far beyond the duplicated locus itself (Belyaev et al. 2004; Bruce et al. 2004; Shimojo 2008). The persistence of these divergent copies across species is consistent with the integration of duplicated genes into distinct regulatory programs following duplication (Ohno 1970a; Ohno 1970b; Birchler and Yang 2022).

A second major finding is that key antigen-recognition genes in bison show pronounced structural differences. Multiple BOLA class I genes in bison exhibit altered signal peptides or novel N-terminal extensions absent from orthologs in cattle and elk. These regions direct MHC class I molecules to the endoplasmic reticulum, where peptide loading and assembly with antigen-processing occur. Even subtle architectural changes may reshape the immunopeptidome presented to T cells (Babiuk et al. 2007; Negessu 2023; Tampé 2025). Several T cell receptor genes show structural modifications, including duplicated protein domains and altered membrane anchors, which can influence antigen recognition and signaling (Shrestha et al. 2009; Haralambieva et al. 2017). Altogether, these modifications point to bison-specific specialization of antigen presentation and T cell recognition pathways, rather than a simple expansion of immune system genes. Structural modifications in MHC molecules could shift the repertoire of peptide epitopes presented, while T cell receptors may change recognition specificity or activation thresholds, both of which are critical determinants of vaccine-induced protective immunity (Reche and Reinherz 2003; Ovsyannikova et al. 2012; Poland et al. 2018).

Gene expression analyses reinforce this interpretation. Weighted gene co-expression network analysis identified discrete immune-associated modules, including a large module enriched for inflammatory response genes and prominently expressed in immune tissues. The presence of structurally divergent immune genes within these modules indicates that architectural novelty is coupled with active transcriptional deployment. Thus, immune gene divergence in bison is not confined to the sequence space but is embedded within coordinated regulatory programs, supporting the biological relevance of these structural differences.

Although the present study does not directly evaluate vaccine responses, the structural and regulatory differences identified here provide candidate mechanisms that may contribute to known differences in immune responses among bison, cattle, and elk. Structural modifications in MHC molecules could alter antigen presentation, whereas variation in T-cell receptor architecture may influence immune recognition and activation (Babiuk et al. 2007; Russell et al. 2022; Negessu 2023; Tampé 2025). Host genetic variation is also known to contribute to differences in vaccine responses among individuals and populations (Ovsyannikova et al. 2012; Haralambieva et al. 2017; Poland et al. 2018). These observations provide candidate mechanisms linking lineage specific genome evolution to variation in immune responses among ungulates.

Together, these findings indicate that immune system evolution in bison and related ungulates has been shaped not only by sequence divergence within immune genes but also by recurrent structural remodeling of regulatory loci. The repeated duplication and divergence of the REST-NOA1 region demonstrates that some genomic regions are particularly prone to generating evolutionary novelty and highlights recurrent duplication as a mechanism capable of reshaping regulatory and immune system architecture over evolutionary timescales. More broadly, these results illustrate how comparative genomics can reveal the processes underlying lineage-specific adaptations and provide a framework for understanding immune-system diversification in wild and domesticated ungulates.

## Data Access

The chromosome-scale genome assembly, gene annotation, and associated sequencing data generated in this study have been deposited at NCBI BioProject PRJNA1397227. All scripts used for this study are publicly available at https://github.com/ISUgenomics/bison2020. https://dataview.ncbi.nlm.nih.gov/object/PRJNA1397227?reviewer=5icup1qbrk84ns9be9ukotj79s

## Competing Interests

The authors declare that they have no competing financial interests or personal relationships that could have influenced the work reported in this study

## Author Contributions

Rick E. Masonbrink performed comparative genomic analyses, gene annotation, orthology analyses, duplication characterization, manuscript preparation, and project coordination. Sharu Paul Sharma created phylogenies. Viswanathan Satheesh created the WGCNA analysis. Aleksandra Badaczewska-Dawid created synteny analyses. Sivanandan Chudalayandi performed repeats analyses on the duplication. Paolo Boggiato and Ellie Putz contribute to study design and sample collection. David Alt contributed to study design. Andrew Severin advised on assembly and annotation practices. Steven Olsen conceived the project, secured funding, and provided project oversight. All authors read the article critically and approved of the final version.

## Notes

### Competing Interest Statement

The authors have declared no competing interest.

